# Shotgun proteomics analysis of nanoparticle-synthesising *Desulfovibrio alaskensis* in response to platinum and palladium

**DOI:** 10.1101/565036

**Authors:** Michael J. Capeness, Lisa Imrie, Lukas F. Mühlbauer, Thierry Le Bihan, Louise E. Horsfall

## Abstract

Platinum and palladium are much sought-after metals of global critical importance in terms of abundance and availability. At the nano-scale these metals are of even higher value due to their catalytic abilities for industrial applications. *Desulfovibrio alaskensis* is able to capture ionic forms of both of these metals, reduce them, and synthesize elemental nanoparticles. Despite this ability very little is known about the biological pathways involved in the formation of these nanoparticles. Proteomic analysis of *D. alaskensis* in response to platinum and palladium has highlighted those proteins involved in both the reductive pathways and the wider stress-response system. A core set of 13 proteins was found in both treatments and consisted of proteins involved in metal transport and reduction. There were also 7 proteins specific to either platinum or palladium. Over-expression of one of these platinum-specific genes, a NiFe hydrogenase small subunit (Dde_2137), resulted in the formation of larger nanoparticles. This study improves our understanding of the pathways involved in the metal resistance mechanism of *Desulfovibrio* and informs how we can tailor the bacterium for nanoparticle production, enhancing its application as a bioremediation tool and as way to capture contaminant metals from the environment.

**Importance:** Bacteria, in particularly *D. alaskensis*, represent a biological and greener way to capture high value metals such as platinum group metals from environmental and industrial waste streams. The recovery of these metals in nanoparticle forms adds extra value to this process as they can be used in a variety of different industrial applications as they have exceptional catalytic capabilities. *D. alaskensis* ability to do this, has been widely reported, though very little is understood about the underlying protein and genetic components. It is by understanding the biological basis of this capability that we can further improve and adapt this bacterium to be better at bioremediation and to control its ability to do so.

## Introduction

There is an increasing demand for expensive platinum group metals. Amongst other applications, they are used for automotive catalytic converters from which platinum and palladium escape via the exhaust and are therefore a source of environmental pollution (1). Although heavy metal ions are toxic to most bacteria, highly-resistant strains are available that absorb them and reduce them to the metallic state. This results in the formation of nanoparticles that can be harvested to provide a source of these valuable metals. Furthermore, small nanoparticles from bacteria have important biotechnological applications, for example, as chemical catalysts. Not only does biological recovery from environmental sources offer an attractive way to recover them from dilute solutions, but also it provides a mechanism for decontaminating polluted environments. Prokaryotes have been used to remediate metal ions from solution for decades: mostly high value metals such as gold, platinum and silver have been targeted (2, 3).

The potential uses of nanoparticles are expanding, from cancer treatment using platinum or arsenic nanoparticles (4, 5) to the use of silver nanoparticles as antimicrobials (6), palladium nanoparticles as catalysts in fuel cells (7) and to treat water contaminated with pharmaceuticals (8). Tailoring these nanoparticles to their implementation for industrial or medical use is vital but so far there has been little standardisation with regards to the size of nanoparticles formed (9). As different sized nanoparticles have different catalytic properties, their use has been limited. Bacteria potentially represent a genetically programmable method for the customisation of nanoparticle size and morphology for specific purposes. As well as this bioremediation by bacteria involves mild conditions and ambient temperatures, thus avoiding the disadvantages of chemical recovery that requires the use of high temperatures or hazardous chemicals.

Anaerobic, sulphate reducing bacteria of the genus *Desulfovibrio* are amongst the most widely used bacteria to remediate metal ions. They reduce multi-valent metal ions to zero-valent or bi-elemental nanoparticles. The list of known metals that it can process is ever increasing and currently consists of chromium, magnesium, iron, technetium, uranium, nickel (10–15), and also includes platinum and palladium: palladium has been the most studied. The *Desulfovibrio* sp. are ideally suited for further analysis as they are already highly tolerant to platinum and palladium ions, and can survive up to 2 and 5 mM respectively (15). They reduce these metals at incredible speed, with up to 100% of Pd^2+^ being converted to Pd(0) within 5 minutes (7).

Only fragmentary information is available at the genetic level concerning *Desulfovibrio* genes required for heavy metal ion reduction. One exception is *cycA* from *D. vulgaris* Hildenborough, which encodes a tetraheme cytochrome c3 involved in uranium reduction *in vitro* (16). The NiFe hydrogenases of *D. fructosovorans* have been shown to be involved in technetium reduction (17). They similarly have a role, along with a Fe-containing hydrogenase, in Pd(0) deposition and clustering in both the periplasm (18) and on cellular membranes (19), while more recently have been implicated in having a roll in palladium deposition in *Shewanella* (20). While important for the anaerobic reduction of palladium by *Desulfovibrio* sp., they are not required for the aerobic reduction of palladium by *E. coli* (21). Large gaps remain in our knowledge of the proteins involved in metal ion reduction. As the particles formed differ between growing and non-growing cultures, in many previous studies washed bacteria were incubated in the presence of the target metal in buffer rather than in substrates required for growth. We therefore designed experiments to identify changes in the proteome of *D. alaskensis* strain G20 during incubation with Pt^4+^ or Pd^2+^ solutions under conditions used in previous studies both by ourselves and many other research teams. One of the genes implicated in Pt^4+^ reduction was cloned into an expression vector in a proof of principle experiment designed to demonstrate that over-production of the gene product results in changes in the physical properties of the nanoparticles formed.

## Experimental Details

### Growth of bacterial strains

*D. alaskensis* G20 was purchased from DSMZ (Deutsche Sammlung von Mikroorganismen und Zellkulturen) and grown in Postgate Medium C (PGMC) using lactate as a carbon source (22). Cultures were grown and manipulated in an anaerobic chamber (Don Whitley) at 30°C in an atmosphere of 10% CO_2_, 10% H_2_ in N_2_. For strains carrying the pMO9075 plasmid the medium was supplemented with 100 μg/ml spectinomycin (Cambridge Bioscience).

### Platinum and palladium treatment

Bacteria were grown in Postgate Medium C to an OD_600_ of 1.0 and centrifuged for 10 min at 4000 rpm. The cell pellets were washed three times in 10 mM MOPS (pH 7.0) and re-suspended finally in MOPS. Solutions of PtCl4 or Na2PdCl4 were then added to a final concentration of 2 mM. The cells in the presence of palladium and platinum and an unsupplemented control suspension were incubated anaerobically at 30°C. After 2 h, samples were centrifuged as before and the supernatant was removed. The pellets were either snap-frozen and stored at −80°C or used immediately for TEM analysis.

### Proteomics analysis: trypsin digestion, LC-MS and data analysis

Cells were disrupted in 8 M urea and the total protein was assayed using a Bradford kit (Biorad, UK). One milligram of protein per sample was digested with trypsin as described previously (23). Briefly, samples were diluted to a final concentration of 2 M urea with 25 mM ammonium bicarbonate and 5 mM DTT. After incubation for 30 min at room temperature, iodoacetamide was added to a final concentration of 12.5 mM. Trypsin (10 μg; Worthington) was added and the sample was digested overnight at room temperature. The samples were cleaned with 25 mg of Bond Elut LMS (Agilent Technologies). Peptides eluted with acetonitrile were aliquoted, dried under low pressure and stored at −20°C until analysis. Prior to LC-MS analysis, 4 μg of the samples were reconstituted in 12 μl of loading buffer (0.05% Trifluoroacetic acid in water) and analysed by capillary-HPLC-MS/MS on an on-line system consisting of a 1200 binary HPLC micro-pump system (Agilent Technologies) coupled to a hybrid LTQ-Orbitrap XL instrument (Thermo-Fisher) on a 140 minute gradient.

MS/MS data were analysed using MASCOT Versions 2.4 (Matrix Science Ltd, UK) against the 3258 sequences in the *D. alaskensis* G20 genome from **http://genome.ornl.gov/microbial/ddes** (24). The Mascot search parameters were two missed-cut. Allowances were made for variable methionine oxidation and fixed cysteine carbamidomethylation in all searches. The precursor mass tolerance was fixed at 7 ppm and MS/MS tolerance to 0.4 amu. The significance threshold was set at 0.05 and an additional cut off peptide score of 20. Progenesis (Nonlinear Dynamics, UK) was used for label-free quantitation. Only peptides that were not shared between different proteins were used for quantification. Data for a sub-set of MS/MS peaks with positive charges of 2, 3, or 4 were extracted from each LC-MS run and the median global ion intensities were extracted for normalization. The abundance of each protein was calculated from the sum of the intensities of unique peptides with positive charges of 2, 3, or 4. Because the method of detection can generate a significant amount of near zero measurements for which a standard log transformation is not ideal, the calculated protein abundances were transformed using an ArcSinH function. The within group means were calculated to evaluate the fold change and the ArcSinH transformed data were then used to calculate the p-values using one-way ANOVA. Proteins were considered to be differentially expressed only if at least two peptides were detected with an absolute ratio of at least 1.5-fold more abundant or 0.667 less abundant and a probability of false discovery of p<0.05, as illustrated in **Figure 1**. The mass spectrometry proteomics data have been deposited to the ProteomeXchange Consortium via the PRIDE partner repository with the dataset identifier PXD004457 and 10.6019/PXD004457 (25). The data are also available in the supplemental material. Functions of the proteins highlighted in this study were predicted using a combination of literature searches and cross-referencing with the KEGG (http://www.kegg.jp/kegg/kegg2.html), STRING (http://string-db.org/), Pfam (http://pfam.xfam.org/) and UniProt databases (http://www.uniprot.org/). The presence of platinum group metals in RNA preparations prevented us from using Quantitative Real-Time PCR to confirm at the transcription level changes in expression of genes for the proteins highlighted.

**Figure 1.**
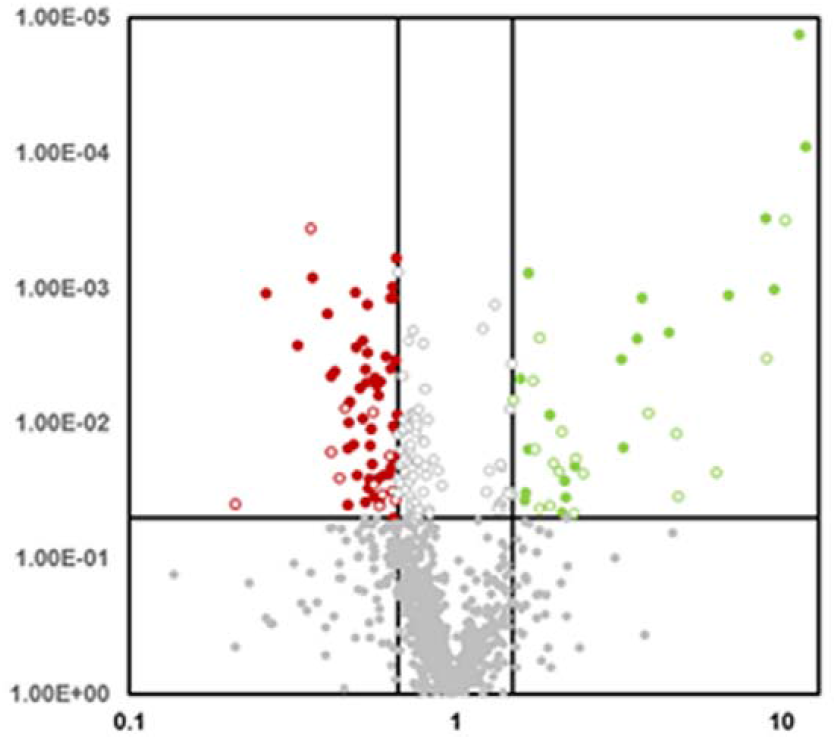
Volcano plot for the data generated in the shotgun proteomics. Red full circles are those proteins less abundant and green full circles are those that are more abundant, both of which are greater than 0.05 probability. Grey circles are proteins that are not significantly different across the three different datasets. Empty circles are those that are significant but are represented by a single peptide only.

### Construction of pMO-2137

The gene for the Dde_2137 subunit of the nickel-iron hydrogenase was cloned using a standard Phusion PCR protocol into the expression vector pMO9075 (26).Forward and reverse primers were dde_2137F 5’-CAGTACTCTGCAGATCTGATCCTTTCATTGC-3’ and dde_2137R 5’-TGCATGCCAATTCGTTTCGGTACCGGC-3 ‘. Restrictions sites for *ScaI* and *SphI* are underlined. The PCR product was digested with ScaI and SphI and ligated into pMO9075 using these restriction sites. The sequence was inserted downstream of the promoter of the kanamycin resistance gene in pMO9075, the expression from which has been shown to complement mutations in *Desulfovibrio* sp.

The pMO-2137 and empty pMO9075 plasmids were transformed into *D. alaskensis* made competent using an adaptation of Li and Krumholz’s method (27). Briefly, cultures at an OD_600nm_ of 0.6-0.8 were centrifuged, washed twice in 400 mM sucrose, 1 mM MgCl_2_, and finally resuspended in 1/10 of the original volume of sucrose solution. One microgram of the plasmid was then added. After 30 min on ice, the suspension was electroporated at 1.5 kV. After electroporation 1 ml of PGMC broth was added immediately to allow the cells to recover at 30°C. After 4 hours, cells were plated onto PGMC agar containing 100 μg/ml of spectinomycin. After 7 days at 30°C, transformants were purified and screened by plasmid isolation for the presence of the construct. For comparative analysis of nanoparticles produced by bacteria transformed with either the empty plasmid control or the plasmid carrying the extra copy of the dde_2137 gene, the OD_600nm_ of the cells were matched before the addition of platinum ions.

### Transmission electron microscopy (TEM)

Samples were drop cast onto a 200 mesh copper grid with a formvar coating (TAAB). The sample was allowed to settle for 5 minutes before excess liquid was removed using filter paper. The grid was then loaded into a JEOL JEM-1400 Plus electron microscope and imaged using a GATAN OneView Camera. The images were processed and the size of the nanoparticles was determined using ImageJ software (http://imagej.nih.gov/ij/).

### Energy-dispersive X-ray spectroscopy

Samples prepared as above for conventional TEM were drop cast onto holey-carbon nickel grids (Agar Scientific). The samples were analysed using a JOEL JEM 2011 TEM fitted with an ISIS system and viewed at an accelerating voltage of 200 kV.

## Results and Discussion

### Overview of the proteomic data

Transfer of a bacterial culture from a growth medium into buffer containing toxic levels of a heavy metal results in an almost total cessation of growth. Amongst many other consequences, this results in ribosome degradation and therefore decreased demand for stable RNA and ribosomal proteins (**Table 1**). The degradation products can then be recycled for the synthesis of components beneficial to the stress response. Conversely, incubation in a buffer in the absence of growth nutrients inevitably limits the ability of bacteria to respond to stress. Any increased protein synthesis is therefore likely to be limited to synthesis of those proteins that increase survival of the stress imposed. These were expected to be proteins involved in the reduction of the specific metal ions, in transporting them out of the cell, or in a more general response to stress. A total of 1061 proteins from the *D. alaskensis* G20 potential proteome were identified by shotgun label-free proteomics, representing 32% of the global proteome (See supplemental material for complete list). In terms of coverage this is in with good keeping with other proteomic studies with metal trails in bacteria, including those of *Desulfovibrio* sp. (28, 29).

**Table 1.**
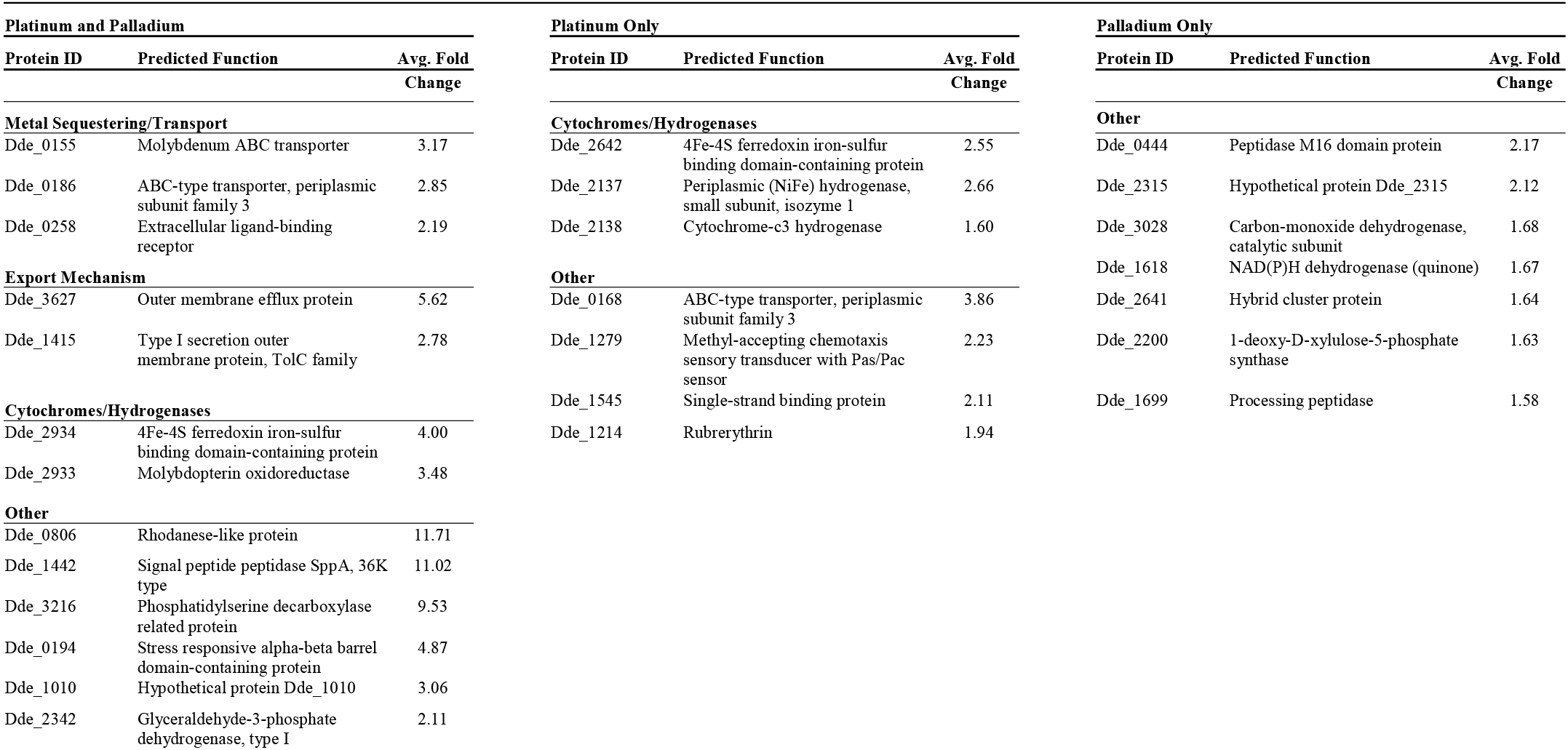
Proteins significantly more abundant (>1.5 fold) in the presence of platinum and palladium and their overlap between datasets.

Using a significance threshold of 1.5-fold change and a p-value below 0.05, the abundance of 46 proteins decreased during incubation in the presence of palladium and platinum ions of which 24 were in the platinum dataset, 22 were in the palladium dataset, with 20 proteins found in both datasets (**Figure 1**). Only 20 proteins were found to be more abundant in cells treated with palladium compared with untreated cells. Similarly, 20 proteins were also more abundant in cells treated with platinum, thirteen of which were also induced during incubation with palladium ions. Consequently, 7 of the more abundant proteins were unique to each dataset (**Figure 2**).

**Figure 2.**
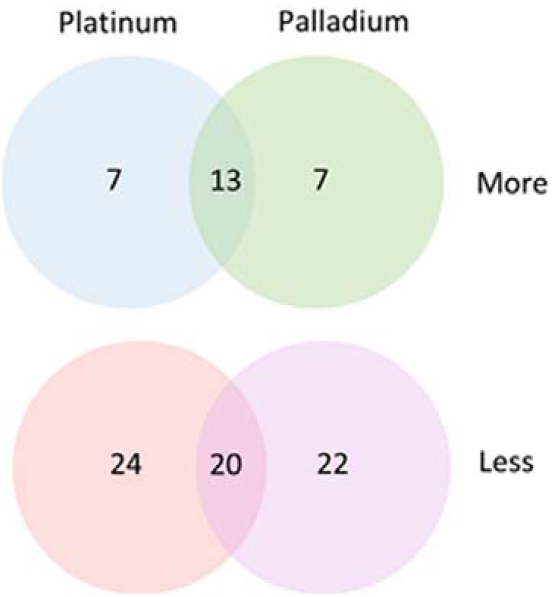
A Venn Diagram showing the number of more/less abundant proteins in the proteomic study and there overlap between the two datasets.

**Figure 3.**
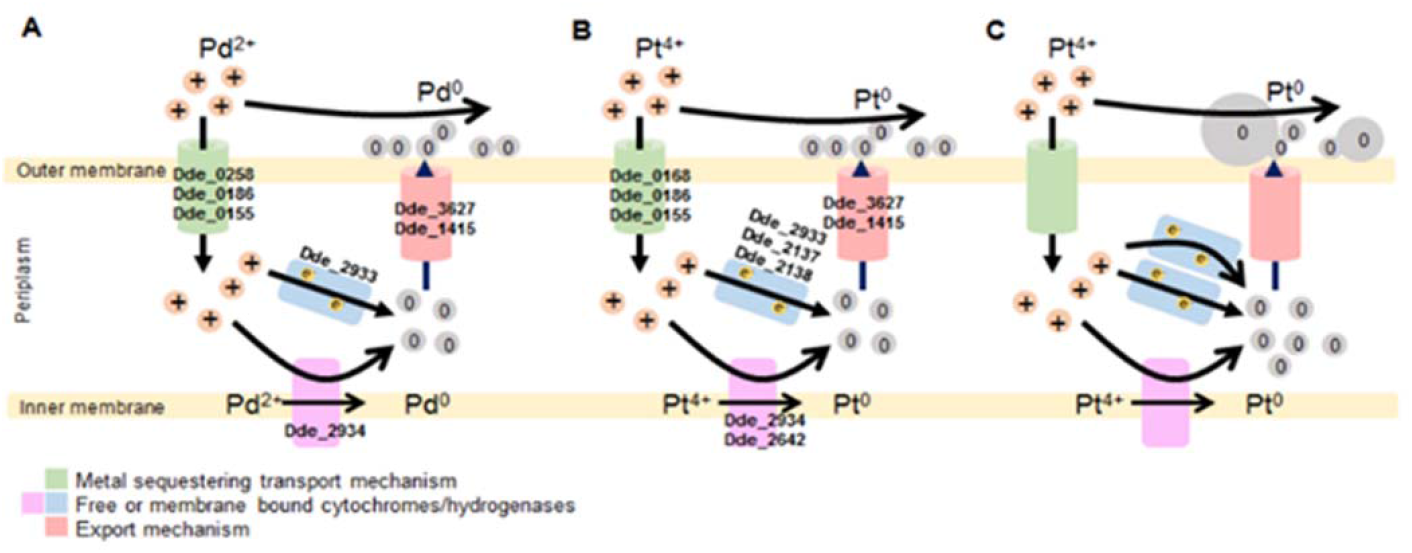
Proposed mechanism of Pd and Pt nanoparticles synthesis in *D. alaskensis* G20 based on the proteomic data presented here for Palladium (**A**) and Platinum (**B**). Note the overlapping of proteins found in each of the individual metal datasets found in the ESI (Table 1). (**C**) Predicted production of increased size platinum nanoparticles due to increased levels of Dde_2137. Adapted and expanded from Capeness *et al*; 2015.

### Proteins less abundant in both the platinum and palladium datasets

As expected, proteins found in lower amounts in both datasets are predominantly those involved in RNA transcription and translation. These include both large and small ribosomal subunit proteins (L32, L29, S187, S12) as well as transcription regulators of the TraR/DskA and MucR families (**Table 2**). One hypothetical protein identified (Dde_0492) is also predicted to be involved in valyl-tRNA synthesis while another (Dde_1811), of unknown function, is only conserved among the *Desulfovibrio* sp. and SRB deltaproteobacteria. The least abundant protein relative to the control was the periplasmic Fe hydrogenase small subunit (Dde_2280), which together with Dde_2274 are potential hydrogenases not involved in the reduction of either platinum or palladium. This suggests that there might be a separate hydrogenase and other electron transfer proteins that are specific for platinum and palladium reduction. Also in the less-abundant dataset are proteins involved in energy production and metabolism such as oxidoreductases (Dde_0584, Dde_1113, Dde_3240, Dde_2272 and Dde_1638), one ATPase subunit (Dde_0990) and the prokaryotic molybdopterin-containing oxidoreductase (Dde_2274).

**Table 2:**
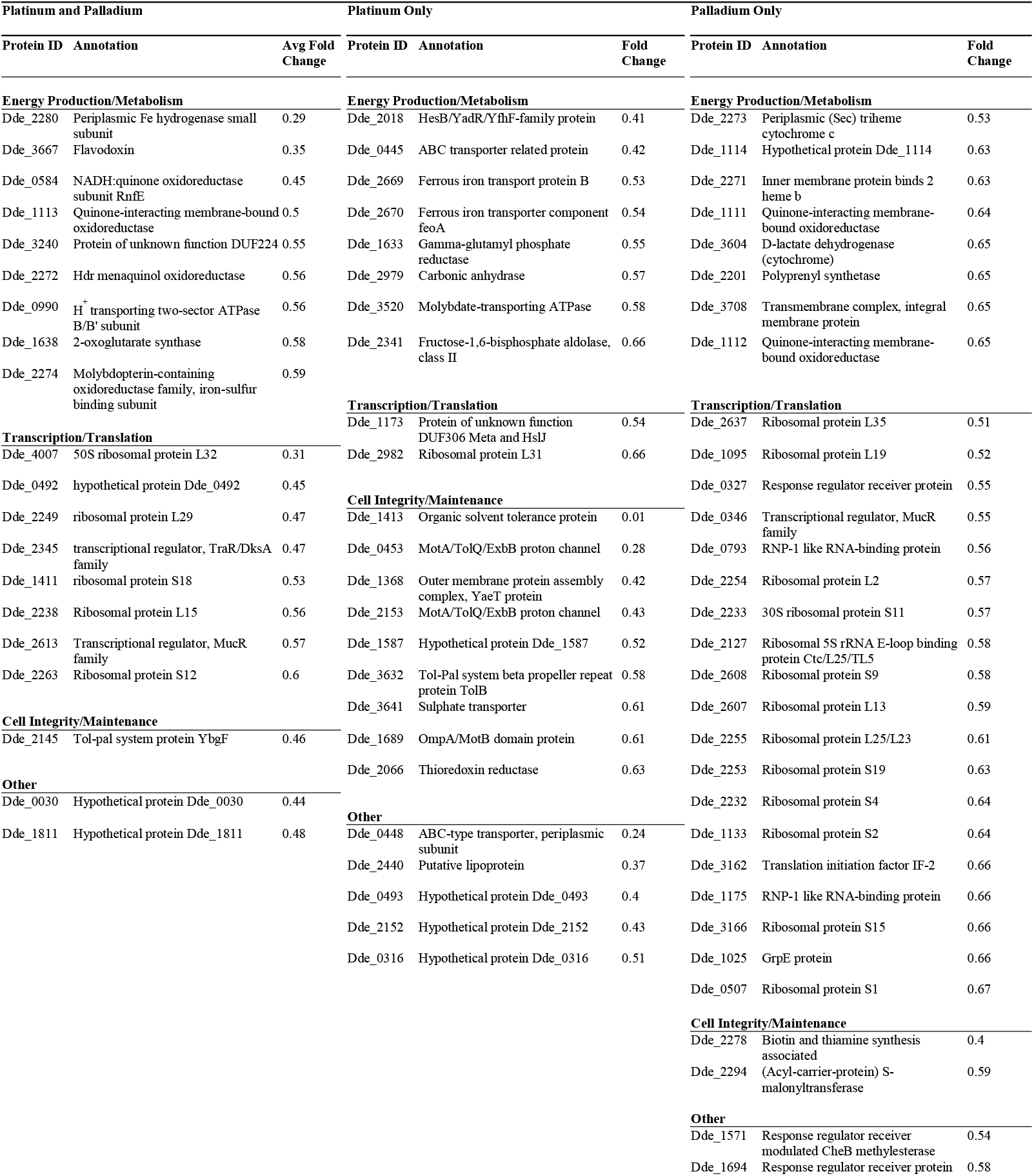
Proteins significantly less abundant (<0.667 fold) in the presence of either platinum and palladium and their overlap between datasets.

### Platinum specific proteins

Proteins involved in cell integrity and maintenance as well as those required for growth dominated the list that were less abundant following incubation with Pt^4+^ ions. Proteins such as Dde_1413, Dde_0453, Dde_1368, Dde_2153, Dde_3632, Dde_1689 and Dde_1131 are predicted to be involved in membrane biogenesis and integrity, which is consistent with decreased synthesis of new membranes due to cessation of growth. This list also includes other proteins with a common role: Dde_3488, Dde_3490, Dde_3567, Dde_2763, Dde_3070, Dde_3703 and Dde_2744 are all predicted to be involved in amino acid synthesis. Again this supports that due to the limited media in which the NPs are produced (MOPS, (3-(N-morpholino)propanesulfonic acid) buffer) there is limited new synthesis of new proteins. Other less abundant proteins Dde_2018, Dde_0445, Dde_2669 and Dde_2670 are required for iron or haem uptake, possibly as a risk reduction strategy due to their general metal binding and transportation ability.

### Palladium specific proteins

The palladium dataset is dominated by ribosomal subunit proteins both small (Dde_2233, Dde_2608, Dde_2253, Dde_2232, Dde_1133) and large (Dde_2637, Dde_1095, Dde_2254, Dde_2607, Dde_2255). There are also other proteins involved in transcription and translation such as the rRNA E-loop binding protein (Dde_2127), the response regulator and CheY-like protein (Dde_0327), the translation initiation factor (Dde_3162) and the MucR transcriptional regulator (Dde_0346). Decreased levels of these proteins would have a global effect on transcription and translation and silencing their expression would arrest the cell cycle. The next largest group that are less abundant specifically in Pd^2+^-treated bacteria are proteins required to provide metabolites for energy production and growth. The proteins Dde_1111 to 1114 (with Dde_1113 found down regulated following both platinum and palladium specific treatment) are products of a group of contiguous genes encoding part of the membrane bound Qmo electron transport complex. The operon is vital to the growth of *D. vulgaris* Hildenborough on lactate-sulphate media (26, 30). Down-regulation of this operon, would make sense as the cells have gone from a lactate-sulphate containing growth medium to a very minimal one.

### Proteins more abundant following incubation with Pt^4+^ or Pd^2+^ ions

Three of the proteins of increased abundance in both datasets included a ferredoxin and a molybdoprotein, MopB, encoded by adjacent genes on the chromosome, and the rhodanese Dde_0806. Evidence that these proteins are required for metal ion reduction includes a report that a *mopB* mutant is unable to grow with hydrogen or formate as the electron donor (31), and that a rhodanese mutant is also unable to use hydrogen as an electron donor (32). More recently it was shown that molybdoproteins are required for Pd^4+^ reduction by *E. coli* (21).

Five other proteins encode putative ABC-type transportation proteins, Dde_0155, Dde_0186, Dde_0258, and an efflux pump, Dde_3627 and Dde_1010, which potentially encodes an outer membrane porin. Also in this group is a member of the TolC family of type I secretion systems (Dde_1415) (**Table 1**). The remaining five proteins common to both datasets all play more general roles in the response to stress. They include Dde_0194, which is an alpha-beta barrel protein, and Dde_1442, an ATP-dependent peptidase likely to be involved in the degradation of proteins that have misfolded due to the presence of the heavy metals (33). Also in this group is Dde_3216, a protein predicted to be involved in lipid metabolism and repair: such proteins would have a role in managing the effects heavy metals or nanoparticles have on lipid membranes. Lipid peroxidation in particular is one of the ways in which bacterial treatment with TiO_2_ nanoparticles (33) and copper ions/nanoparticles act as antimicrobials (34). Finally Dde_2342 is glyceraldehyde-3-phosphate dehydrogenase, which understandably would play a critical role in glycolysis as the cells metabolise endogenous reserves for ATP production.

### Proteins accumulated only in response to exposure to Pt^4+^ ions

Some proteins accumulated in the presence of Pt ions but not Pd ions and vice versa. Included in the platinum-specific dataset are Dde_2137, which is the small subunit of the NiFe hydrogenase, and Dde_2138, which a c_3_-type cytochrome that is believed to donate electrons to the NiFe hydrogenase active site. The NiFe hydrogenase subunit is transported into the periplasm by the twin arginine targeting TAT pathway. Both of these proteins were shown to be required for metal ion reduction by *D. fructosovorans* (17). The NiFe hydrogenase genes are flanked by two genes encoding the large and small subunits of an alternative NiFeSe hydrogenase and maturation proteins, but none of these other proteins were more abundant following incubation with Pt^4+^. Another ferredoxin, Dde_2642, also accumulated during incubation with Pt^4+^ ions. This protein is thought to be involved in metal-binding and electron transfer in the cytoplasm. Whether it is able to shuttle electrons to the platinum ions in *Desulfovibrio* is unknown.

An outer membrane efflux protein, Dde_3267, accumulated only in the platinum-treated bacteria. This could be involved either in the export of platinum ions out of the cell to limit their cytotoxic effect, or even in the efflux of the nanoparticle itself once it has been formed.

The ability of platinum ions to cause DNA lesions that inhibit cell division is the basis for their chemotherapeutic use in drugs such as cisplatin. The single strand binding (SSB) protein (Dde_1545) accumulated in bacteria incubated with Pt^4+^ ions (35). Binding of Pt is prevented by the binding of SSB to single stranded DNA. Two other potential stress response proteins accumulated during incubation with Pt^4+^ ions. One was rubrerythrin (Dde_1214) that due to its ability to protect *D. vulgaris* against hydrogen peroxide damage is suggested to protect against cytoplasmic oxidative stress (36, 37), it was also previously found to be down-regulated when using nitrate as electron acceptor (38). The other is a methyl-accepting chemotaxis sensory transducer (Dde_1279), which contains a PAS domain at the conserved-C-terminus that is likely to be part of a signal response to promote motility in the presence of platinum.

### Proteins accumulated only in response to exposure to Pd^2+^ ions

No obvious candidates for redox proteins required for Pd^2+^ reduction accumulated specifically in response to incubation with Pd^2+^ ions. Although carbon monoxide reductase (Dde_3028), the hybrid cluster protein (Hcp) (Dde_2640) and a quinone-linked NAD(P)H dehydrogenase (Dde_1618) were more abundant than in the control, these increases were just over 1.5-fold. The Hcp in particular has been implicated in response to nitric oxide stress in both *D. vulgaris* (39) and *E. coli* (40). The four other proteins that increased significantly included a peptidase that might be involved in a stress response, but again the increases relative to the control were only about two-fold. The Dde_2200 protein is a 1-deoxy-D-xylulose-5-phosphate synthase involved in the metabolism of pyruvate, a key intermediate in various metabolic pathways. Since the cells are in buffer (with palladium), it is likely that this would increase metabolism of what little pyruvate is available. Similarly Dde_1699 and Dde_0444, both of which are zinc-containing peptidases, are also present and are likely to be degrading proteins that have misfolded due to the presence of the palladium/oxidative stress involved in metal reduction (41). Clearly, however, most of the proteins induced in response to exposure to Pd^2+^ were those also induced in the presence of Pt^4+^ ions.

### Increase in nanoparticle sizes due to increased synthesis of the NiFe hydrogenase

A major aim of the proteomics experiments was to investigate whether proteins could be identified that when over-expressed might facilitate the controlled production of metal nanoparticles of a specific size. Given the previous demonstration that a mutant defective in NiFe hydrogenase synthesis was also defective in nanoparticle production, the observation that this enzyme accumulated during Pt^4+^ reduction was especially significant. We therefore designed experiments to determine whether over-expression of the NiFe hydrogenase would result in changes in the physical properties of the nanoparticles formed. The gene encoding the NiFe hydrogenase protein Dde_2137 was cloned into the *D. alaskensis* plasmid pMO9075 to give pMO-2137. Introduction of this plasmid to *D. alaskensis* G20 significantly increased the size of the Pt nanoparticles when cells were exposed to platinum ions (p<<0.005) (**Figure 4**). On average the size increased from 117 nmfor the empty plasmid control to 324 nm for the strain carrying the pMO-2137. Increased production of the NiFe protein also increased the range of nanoparticle sizes formed (**Figure 5** and **Figure 6**). Many of the larger particles were free in the buffer, not attached to the cell. Though larger nanoparticles were formed, the transformants were not more resistant to platinum treatment. We therefore suggest that the increased amount of Dde_2137 leads to increased deposition of platinum nanoparticles on the cell surface. This leads to an increased deposition rates and, in turn, an increased localised turnover rate of the platinum ions by the autocatalytic abilities of the platinum nanoparticles thus leading overall to larger nanoparticles. This supports previous work which concluded that the NiFe hydrogenase is required for nanoparticle synthesis in *Desulfovibrio* (18) and also their deposition on the outer membranes in *Shewanella* (20). Samples of nanoparticles produced by *Desulfovibrio* transformed with either pMO-2137 or the empty vctor control were analysed using EDS (Energy-dispersive spectroscopy). In both cases the nanoparticles were shown to be entirely made of platinum with some regions containing very regular atomic lattices (**Figure 7**).

**Figure 4.**
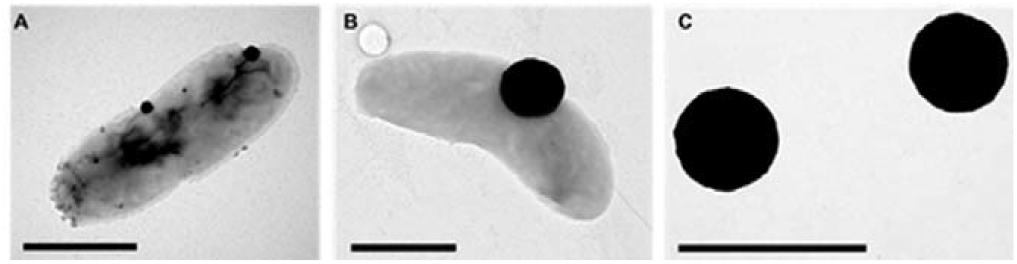
Electron micrographs of *D. alaskensis* incubated with 2 mM PtCl_4_ for 120 minutes. **A**. Control containing the empty pMO9075 plasmid. **B**. containing the pMO-2137 plasmid. **C**. The nanoparticles from B. free in the media. Scale bar = 1 μm

**Figure 5.**
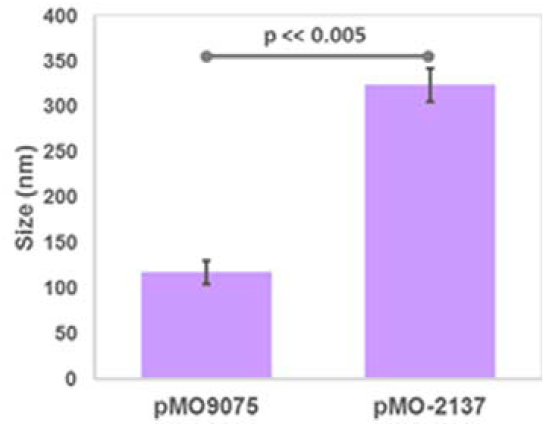
Size distribution of the platinum nanoparticles produced by *D. alaskensis* containing the two different plasmids. The pM09075 control and pM09075 containing gene encoding Dde_2137. Error bars represent the standard error around the mean.

**Figure 6.**
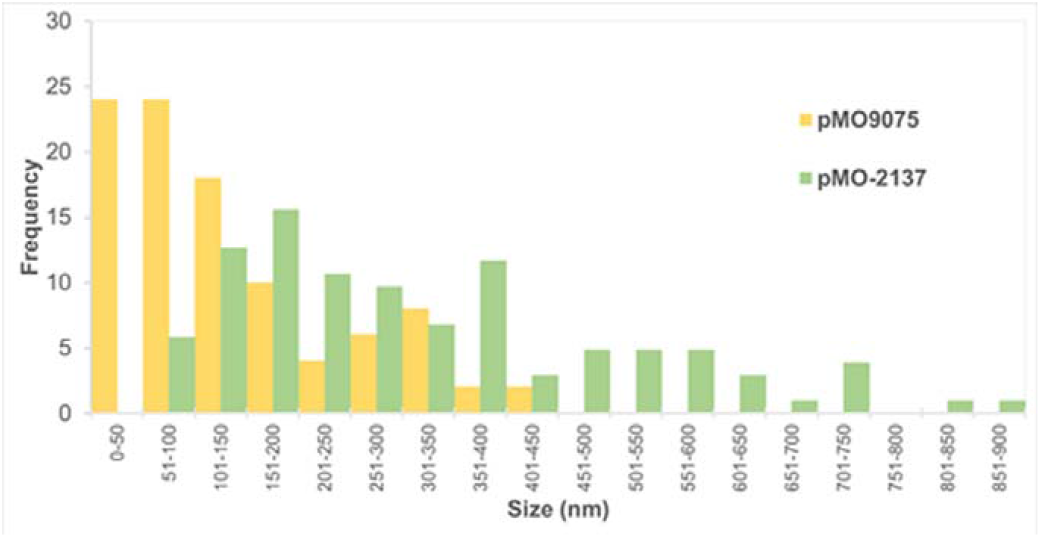
Frequency distribution of the sizes of the platinum nanoparticles produced by *D. alaskensis* containing either the control plasmid (pMO9075) or the one containing the gene dde_2137 encoding the small subunit of the NiFe hydrogenase (pMO-2137).

**Figure 7.**
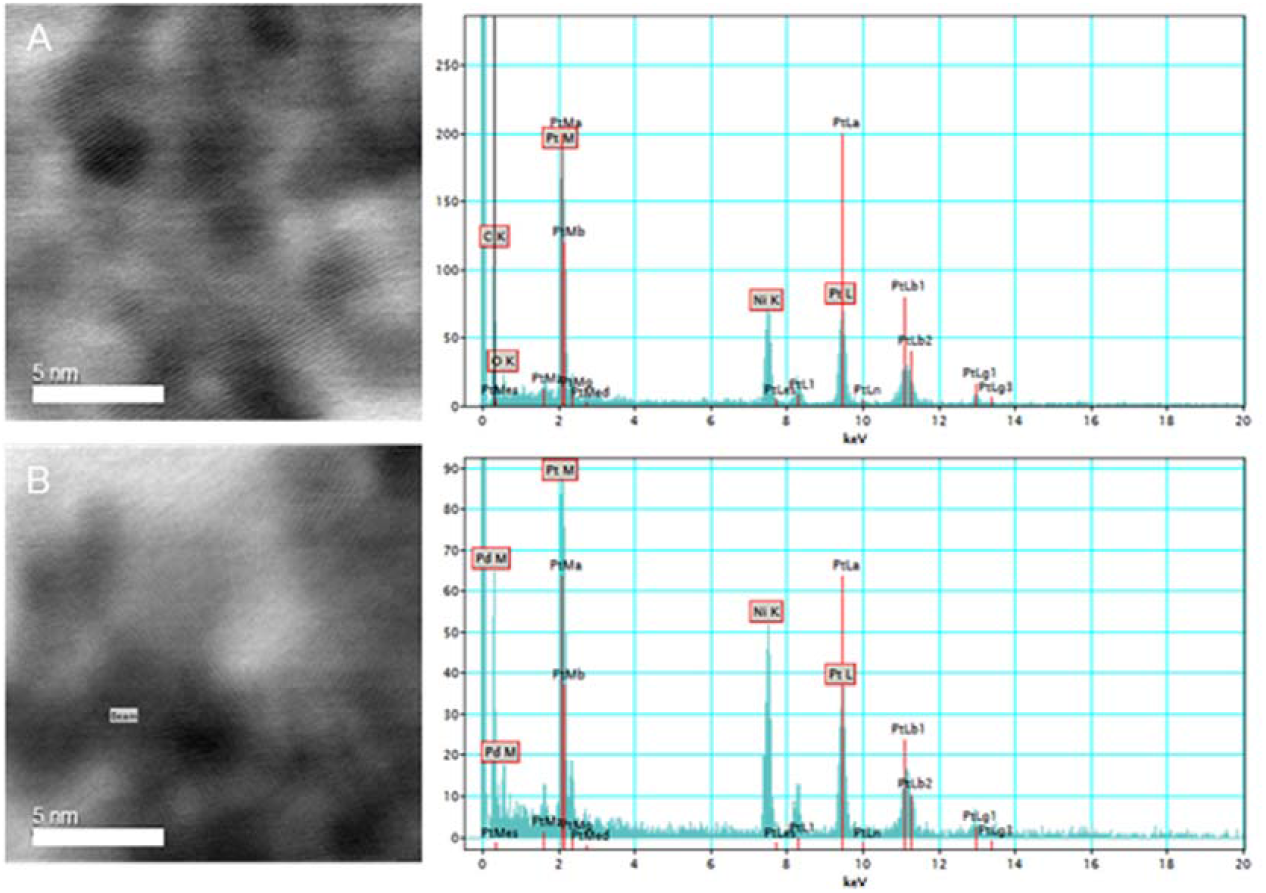
Samples of nanoparticles produced by *D. alaskensis* cells in the presence of platinum. (**A**) Cells containing the empty plasmid pMO9075 and attributed EDS spectra confirming the presence of only platinum, (**B**) cells containing the pMO-2137 plasmid expressing the small NiFe subunit, as before, the associated EDS spectrum shows only the presence of platinum. Note: the presence of nickel in the sample is due to the samples being mounted on a nickel grid.

## Conclusions

Despite years of use of *D. alaskensis* for metal ion bioremediation, this paper to our knowledge presents the first shotgun proteomic analysis of bacteria synthesizing palladium and platinum nanoparticles. Although various strains of *Desulfovibrio* were used for these previous studies, only a few of the proteins reported here have previously been implicated in metal reduction and nanoparticle production.

We have identified some proteins that accumulate during both palladium or platinum reduction and others that respond to one of the metals, but not the other. This implies that there are separate pathways for reduction of each metal, but also an underlying core pathway that features in the reduction of both. One possible reason for this is that Pt^4+^ ions are reduced via Pt^2+^ and that the divalent Pd^2+^ and Pt^2+^ cations are reduced by a common pathway. Identification of both groups of these proteins presents the opportunity not only of increasing the removal of metals from platinum and palladium-containing solutions, but also for increasing the specificity of the process. Genetic manipulation to control the expression of the genes for these target proteins might enable increase metal specificity, efficiency and even metal resistance to be achieved. In proof of principle experiments to test these ideas, we have also shown the feasibility of genetically engineering *D. alaskensis* to alter nanoparticle sizes. One aim of future work will be to develop the ability to produce nanoparticles tailored for specific needs as well as the application of what we have engineered to environmental samples.

## Conflicts of interest

There are no conflicts of interest to declare.

## Acknowledgements

This work is funded by EPSRC (EP/K026216/1 and EP/N026519/1). We would like to thank S. Mitchell for his help with the electron microscopy as part of the Wellcome Trust Multi User Equipment Grant (WT104915MA). We would also like to thank Dr. Sam McFadzean at the University of Glasgow.

## References

1. Gordon RB, Bertram M, Graedel TE. 2006. Metal stocks and sustainability. Proc Natl Acad Sci U S A 103:1209–1214.

2. Pantidos N, Horsfall LE. 2014. Biological Synthesis of Metallic Nanoparticles by Bacteria, Fungi and Plants. J Nanomed Nanotechnol 5.

3. Hulkoti NI, Taranath TC. 2014. Biosynthesis of nanoparticles using microbes-a review. Colloids Surf B Biointerfaces 121:474–483.

4. Porcel E, Liehn S, Remita H, Usami N, Kobayashi K, Furusawa Y, Le Sech C, Lacombe S. 2010. Platinum nanoparticles: a promising material for future cancer therapy? Nanotechnology 21:85103.

5. Balaz P, Sedlak J, Pastorek M, Cholujova D, Vignarooban K, Bhosle S, Boolchand P, Bujnakova Z, Dutkova E, Kartachova O, Stalder B. 2012. Arsenic sulphide As4S4 nanoparticles: Physico-chemical properties and anticancer effects. Journal of Nano Research 18–19:149–155.

6. Gong P, Li HM, He XX, Wang KM, Hu JB, Tan WH, Zhang SC, Yang XH. 2007. Preparation and antibacterial activity of Fe3O4@Ag nanoparticles. Nanotechnology 18.

7. Yong P, Rowson NA, Farr JP, Harris IR, Macaskie LE. 2002. Bioreduction and biocrystallization of palladium by *Desulfovibrio desulfuricans* NCIMB 8307. Biotechnol Bioeng 80:369–379.

8. Martins M, Mourato C, Sanches S, Noronha JP, Crespo MT, Pereira IA. 2017. Biogenic platinum and palladium nanoparticles as new catalysts for the removal of pharmaceutical compounds. Water Res 108:160–168.

9. Delay M, Frimmel FH. 2012. Nanoparticles in aquatic systems. Anal Bioanal Chem 402:583–592.

10. Lovley DR, Phillips EJ. 1992. Reduction of uranium by *Desulfovibrio desulfuricans*. Appl Environ Microbiol 58:850–856.

11. Lloyd JR, Yong P, Macaskie LE. 1998. Enzymatic recovery of elemental palladium by using sulfate-reducing bacteria. Appl Environ Microbiol 64:4607–4609.

12. Lloyd JR, Ridley J, Khizniak T, Lyalikova NN, Macaskie LE. 1999. Reduction of technetium by Desulfovibrio desulfuricans: biocatalyst characterization and use in a flowthrough bioreactor. Appl Environ Microbiol 65:2691–2696.

13. Chardin B, Dolla A, Chaspoul F, Fardeau ML, Gallice P, Bruschi M. 2002. Bioremediation of chromate: thermodynamic analysis of the effects of Cr(VI) on sulfate-reducing bacteria. Appl Microbiol Biotechnol 60:352–360.

14. Payne RB, Gentry DM, Rapp-Giles BJ, Casalot L, Wall JD. 2002. Uranium reduction by *Desulfovibrio desulfuricans* strain G20 and a cytochrome c_3_ mutant. Appl Environ Microbiol 68:3129–3132.

15. Capeness MJ, Edmundson MC, Horsfall LE. 2015. Nickel and platinum group metal nanoparticle production by *Desulfovibrio alaskensis G20*. N Biotechnol 32:727–731.

16. Lovley DR, Widman PK, Woodward JC, Phillips EJ. 1993. Reduction of uranium by cytochrome c_3_ of *Desulfovibrio vulgaris*. Appl Environ Microbiol 59:3572–3576.

17. De Luca G, de Philip P, Dermoun Z, Rousset M, Vermeglio A. 2001. Reduction of technetium(VII) by *Desulfovibrio fructosovorans* is mediated by the nickel-iron hydrogenase. Appl Environ Microbiol 67:4583–4587.

18. Mikheenko IP, Rousset M, Dementin S, Macaskie LE. 2008. Bioaccumulation of palladium by *Desulfovibrio fructosivorans* wild-type and hydrogenase-deficient strains. Appl Environ Microbiol 74:6144–6146.

19. Deplanche K, Woods RD, Mikheenko IP, Sockett RE, Macaskie LE. 2008. Manufacture of stable palladium and gold nanoparticles on native and genetically engineered flagella scaffolds. Biotechnol Bioeng 101:873–880.

20. Dundas CM, Graham AJ, Romanovicz DK, Keitz BK. 2018. Extracellular Electron Transfer by *Shewanella oneidensis* Controls Palladium Nanoparticle Phenotype. ACS Synth Biol doi:10.1021/acssynbio.8b00218.

21. Foulkes JM, Deplanche K, Sargent F, Macaskie LE, Lloyd JR. 2016. A Novel Aerobic Mechanism for Reductive Palladium Biomineralization and Recovery by *Escherichia coli*. Geomicrobiology Journal 33:230–236.

22. Postgate JR, Kent HM, Robson RL, Chesshyre JA. 1984. The genomes of Desulfovibrio gigas and D. vulgaris. J Gen Microbiol 130:1597–1601.

23. Le Bihan T, Grima R, Martin S, Forster T, Le Bihan Y. 2010. Quantitative analysis of low-abundance peptides in HeLa cell cytoplasm by targeted liquid chromatography/mass spectrometry and stable isotope dilution: emphasising the distinction between peptide detection and peptide identification. Rapid Commun Mass Spectrom 24:1093–1104.

24. Hauser LJ, Land ML, Brown SD, Larimer F, Keller KL, Rapp-Giles BJ, Price MN, Lin M, Bruce DC, Detter JC, Tapia R, Han CS, Goodwin LA, Cheng JF, Pitluck S, Copeland A, Lucas S, Nolan M, Lapidus AL, Palumbo AV, Wall JD. 2011. Complete Genome Sequence and Updated Annotation of *Desulfovibrio alaskensis* G20. Journal of Bacteriology 193:4268–4269.

25. Vizcaino JA, Csordas A, del-Toro N, Dianes JA, Griss J, Lavidas I, Mayer G, Perez-Riverol Y, Reisinger F, Ternent T, Xu QW, Wang R, Hermjakob H. 2016. 2016 update of the PRIDE database and its related tools (vol 44, pg D447, 2016). Nucleic Acids Research 44:11033–11033.

26. Keller KL, Rapp-Giles BJ, Semkiw ES, Porat I, Brown SD, Wall JD. 2014. New model for electron flow for sulfate reduction in *Desulfovibrio alaskensis G20*. Appl Environ Microbiol 80:855–868.

27. Li X, Krumholz LR. 2007. Regulation of arsenate resistance in *Desulfovibrio desulfuricans* G20 by an arsRBCC operon and an arsC gene. J Bacteriol 189:3705–3711.

28. Queiroz PS, Ruas FAD, Barboza NR, Borges WD, Guerra-Sa R. 2018. Alterations in the proteomic composition of Serratia marcescens in response to manganese (II). Bmc Biotechnology 18.

29. Qian C, Chen HM, Johs A, Lu X, An J, Pierce EM, Parks JM, Elias DA, Hettich RL, Gu BH. 2018. Quantitative Proteomic Analysis of Biological Processes and Responses of the Bacterium Desulfovibrio desulfuricans ND132 upon Deletion of Its Mercury Methylation Genes. Proteomics 18.

30. Zane GM, Yen HC, Wall JD. 2010. Effect of the deletion of qmoABC and the promoter-distal gene encoding a hypothetical protein on sulfate reduction in *Desulfovibrio vulgaris* Hildenborough. Appl Environ Microbiol 76:5500–5509.

31. Li XZ, Luo QW, Wofford NQ, Keller KL, McInerney MJ, Wall JD, Krumholz LR. 2009. A Molybdopterin Oxidoreductase Is Involved in H2 Oxidation in *Desulfovibrio desulfuricans* G20. Journal of Bacteriology 191:2675–2682.

32. Krumholz LR, Bradstock P, Sheik CS, Diao YW, Gazioglu O, Gorby Y, McInerney MJ. 2015. Syntrophic Growth of *Desulfovibrio alaskensis* Requires Genes for H-2 and Formate Metabolism as Well as Those for Flagellum and Biofilm Formation. Applied and Environmental Microbiology 81:2339–2348.

33. Kubacka A, Diez MS, Rojo D, Bargiela R, Ciordia S, Zapico I, Albar JP, Barbas C, Martins dos Santos VA, Fernandez-Garcia M, Ferrer M. 2014. Understanding the antimicrobial mechanism of TiO(2)-based nanocomposite films in a pathogenic bacterium. Sci Rep 4:4134.

34. Hong R, Kang TY, Michels CA, Gadura N. 2012. Membrane lipid peroxidation in copper alloy-mediated contact killing of *Escherichia coli*. Appl Environ Microbiol 78:1776–1784.

35. Johnstone TC, Alexander SM, Lin W, Lippard SJ. 2014. Effects of monofunctional platinum agents on bacterial growth: a retrospective study. J Am Chem Soc 136:116–118.

36. Pierik AJ, Wolbert RB, Portier GL, Verhagen MF, Hagen WR. 1993. Nigerythrin and rubrerythrin from *Desulfovibrio vulgaris* each contain two mononuclear iron centers and two dinuclear iron clusters. Eur J Biochem 212:237–245.

37. Lumppio HL, Shenvi NV, Summers AO, Voordouw G, Kurtz DM, Jr. 2001. Rubrerythrin and rubredoxin oxidoreductase in *Desulfovibrio vulgaris:* a novel oxidative stress protection system. J Bacteriol 183:101–108.

38. Cadby IT, Faulkner M, Cheneby J, Long J, van Helden J, Dolla A, Cole JA. 2017. Coordinated response of the *Desulfovibrio desulfuricans* 27774 transcriptome to nitrate, nitrite and nitric oxide. Sci Rep 7:16228.

39. Figueiredo MCO, Lobo SAL, Sousa SH, Pereira FP, Wall JD, Nobre LS, Saraiva LM. 2013. Hybrid Cluster Proteins and Flavodiiron Proteins Afford Protection to *Desulfovibrio vulgaris* upon Macrophage Infection. Journal of Bacteriology 195:2684–2690.

40. Wang J, Vine CE, Balasiny BK, Rizk J, Bradley CL, Tinajero-Trejo M, Poole RK, Bergaust LL, Bakken LR, Cole JA. 2016. The roles of the hybrid cluster protein, Hcp and its reductase, Hcr, in high affinity nitric oxide reduction that protects anaerobic cultures of *Escherichia coli* against nitrosative stress. Mol Microbiol 100:877–892.

41. Gur E, Biran D, Ron EZ. 2011. Regulated proteolysis in Gram-negative bacteria--how and when? Nat Rev Microbiol 9:839–848.

